# The recombination landscape in wild house mice inferred using population genomic data

**DOI:** 10.1101/112094

**Authors:** Tom R. Booker, Rob W. Ness, Peter D. Keightley

## Abstract

Characterizing variation in the rate of recombination across the genome is important for understanding many evolutionary processes. The landscape of recombination has been studied previously in the house mouse, *Mus musculus,* and it is known that the different subspecies exhibit different suites of recombination hotspots. However, it is not established whether broad-scale variation in the rate of recombination is conserved between the subspecies. In this study, we construct a fine-scale recombination map for the Eastern house mouse subspecies, *M. m. castaneus,* using 10 individuals sampled from its ancestral range. After inferring phase, we use LDhelmet to construct recombination maps for each autosome. We find that the spatial distribution of recombination rate is strongly positively between our *castaneus* map and a map constructed using inbred lines of mice derived predominantly from *M. m. domesticus.* We also find that levels of genetic diversity in *M. m. castaneus* are positively correlated with the rate of recombination, consistent with pervasive natural selection acting in the genome. Our study suggests that recombination rate variation is conserved at broad scales between *M. musculus subspecies*.

## Introduction

In many species, rates of crossing-over are not uniformly distributed across chromosomes, and understanding this variation and its causes is important for many aspects of molecular evolution. Experiments in laboratory strains or managed populations examining the inheritance of markers through pedigrees have allowed direct estimation of rates of crossing over in different regions of the genome. Studies of this kind are impractical for many wild populations, where pedigree structures are largely unknown (but see Johnston *et al.* 2016). In mice, there have been multiple genetic maps published (e.g. Jensen-Seaman *et al.* 2004; Paigen *et al.* 2008; Cox *et al.* 2009; Liu *et al.* 2014), typically using the classical inbred laboratory strains, which are predominantly derived from the Western European house mouse subspecies, *Mus musculus domesticus* (Yang *et al.* 2011). Recombination rate variation in laboratory strains may not, therefore, reflect natural rates and patterns in wild mice of different subspecies. In addition, recombination rate modifiers may have become fixed in the process of laboratory strain management. On the other hand, directly estimating recombination rates in wild house mice is not feasible without both a population’s pedigree and many genotyped individuals (but see Wang *et al.* 2017).

To understand variation in recombination rates, patterns of linkage disequilibrium (LD) in a sample of individuals drawn from a population can be used. Coalescent-based methods have been developed that use such data to indirectly estimate recombination rates at very fine scales (Hudson 2001; Mcvean *et al.* 2002; Mcvean *et al.* 2004; Auton and Mcvean 2007; Chan *et al.* 2012). The recombination rates estimated in this way reflect variation in crossing over rates in populations ancestral to the extant population, and are averages between the sexes. Methods using LD have been applied to explore variation in recombination rates among mammals and other eukaryotes, and have demonstrated that recombination hotspots are associated with specific genomic features (Myers *et al.* 2010; Paigen and Petkov 2010; Singhal *et al.* 2015).

The underlying mechanisms explaining the locations of recombination events have been the focus of much research. In house mice and in most other mammals, the PRDM9 zinc-finger protein binds to specific DNA motifs, resulting in an increased probability of double-strand breaks, which can then be resolved by reciprocal crossing-over (Grey *et al.* 2011; Baudat *et al.* 2013). Accordingly, it has been shown that recombination hotspots are enriched for PRDM9 binding sites (Myers *et al.* 2010; Brunschwig *et al.* 2012). PRDM9-knockout mice still exhibit hotspots, but in dramatically different genomic regions (Brick *et al.* 2012). Variation in PRDM9, specifically in the exon encoding the zinc-finger array, results in different binding motifs (Baudat *et al.* 2010). Davies *et al.* (2016) generated a line of mice in which the exon encoding the portion of the PRDM9 protein specifying the DNA binding motif was replaced with the orthologous human sequence. The recombination hotspots they observed in this ‘humanized’ line of mice were enriched for the PRDM9 binding motif observed in humans.

Great ape species have different alleles of the PRDM9 gene (Schwartz *et al.* 2014) and relatively little hotspot sharing (Winckler *et al.* 2005; Stevison *et al.* 2015). Correlations between the broad-scale recombination landscapes of the great apes are, however, relatively strongly positive (Stevison *et al.* 2011; Stevison *et al.* 2015). This suggests that, while hotspots evolve rapidly, the overall genetic map changes more slowly. Indeed, multiple closely related species pairs with different hotspot locations show correlations between recombination rates at broad scales (Smukowski and Noor 2011), as do species that share hotspots or lack them altogether (Singhal *et al.* 2015; Smukowski Heil *et al.* 2015).

It has been suggested that a population ancestral to the *M. musculus* species complex began to split into the present day subspecies around 500,000 years ago (Geraldes *et al.* 2008). In this time, functionally distinct alleles of the PRDM9 gene and different suites of hotspots have evolved in the subspecies (Smagulova *et al.* 2016). In addition, between members of the *M. musculus* subspecies complex, there is also variation in recombination rates at relatively broad scales for multiple regions of the genome (Dumont *et al.* 2011), and there is genetic variation in recombination rate is polymorphic between *M. m. domesticus* individuals (Wang *et al.* 2017). Brunschwig *et al.* (2012) analyzed single nucleotide polymorphism (SNP) data for classical laboratory strains of mice, and used an LD-based approach to estimate the sex-averaged recombination landscape for the 19 mouse autosomes. The recombination rate map they constructed is similar to a genetic map generated using crosses by Cox *et al.* (2009). Both studies were conducted using the classical inbred lines (whose ancestry is largely *M. m. domesticus),* and their estimated recombination rate landscapes may therefore reflect that of *M. m. domesticus* more than other members of the *M. musculus* species complex.

In this study, we construct a recombination map for the house mouse subspecies *M. m. castaneus.* We used the genome sequences of 10 wild-caught individuals of *M. m. castaneus* from the species’ expected ancestral range, originally reported by Halligan *et al.* (2013). In our analysis, we first phased SNPs and estimated rates of error in phasing. Secondly, we simulated data to assess the power of estimating recombination rates based on 10 individuals and the extent by which phase errors lead to biased estimates of the rate of recombination. Finally, using an LD-based approach, we inferred a sex-averaged map of recombination rates and compared this to previously published genetic maps for *M. musculus.* We show that variation in recombination rates in *M. m. castaneus* is very similar to rate variation estimated in the classical inbred strains. This suggests that, at broad scales, recombination rates have been relatively highly conserved since the subspecies began to diverge.

## Materials and Methods

### Polymorphism data for Mus musculus castaneus

We analyzed the genomes of 10 wild-caught *M. m. castaneus* individuals sequenced by Halligan *et al.* (2013). Samples were from North-West India, a region that is believed to be within the ancestral range of the house mouse. Mice from this region have among the highest levels of genetic diversity among the *M. musculus* subspecies (Baines and Harr 2007). In addition, the individuals sequenced represent a single population cluster and showed little evidence for substantial inbreeding (Halligan *et al.* 2010). Halligan *et al.* (2013) sequenced individual genomes to high coverage using multiple libraries of Illumina paired-end reads, which were mapped to the mm9 reference genome using BWA (Li and Durbin 2009). Mean coverage was >20x and the proportion of the genome with >10x coverage was more than 80% for all individuals sampled (Halligan *et al.* 2013). Variants were called with the Samtools *mpileup* function (Li *et al.* 2009) using an allele frequency spectrum (AFS) prior. The AFS was obtained by iteratively calling variants until the spectrum converged. After the first iteration, all SNPs at frequencies >0.5 were swapped into the mm9 genome to construct a reference genome for *M. m. castaneus,* which was used for subsequent variant calling (for further details see Halligan *et al.* 2013). The variant call format files generated by Halligan *et al.* (2013) were used in this study. In addition, alignments of *Mus famulus* and *Rattus norvegicus* to the mm9 genome, also generated by Halligan *et al.* (2013), were used as outgroups.

For the purposes of estimating recombination rates, variable sites were filtered on the basis of several conditions: Insertion/deletion polymorphisms were excluded, because the method used to phase variants (*see below)* cannot process these sites. We also excluded sites with more than two alleles and those that failed the Samtools Hardy-Weinberg equilibrium test (*p* < 0.002).

### Inferring phase and estimating switch error rates

LDhelmet estimates recombination rates from a sample of phased chromosomes or haplotypes drawn from a population. To estimate haplotypes, heterozygous SNPs called in *M. m. castaneus* were phased using read-aware phasing in ShapeIt2 (Delaneau *et al.* 2013). ShapeIt2 uses sequencing reads that span multiple heterozygous variants, phase-informative reads (PIRs), and LD to phase variants at the level of whole chromosomes. Incorrectly phased heterozygous SNPs, termed switch errors, may upwardly bias estimates of the recombination rate, because they appear identical to legitimate crossing over events. To assess the impact of incorrect phasing on our recombination rate inferences, we quantified the switch error rate as follows. The population sample of *M. m. castaneus* comprised of seven females and three males. The X-chromosome variants in males therefore represent perfectly phased haplotypes. We merged the BAM alignments of short reads for the X-chromosome of the three males (samples H12, H28 and H34 from Halligan *et al.* (2013)) to make three datasets of pseudo-females, which are female-like, but in which the true haplotypes are known (H12+H28 = H40; H12+H34 = H46; H28 + H34 = H62). We then jointly re-called variants in the seven female samples plus the three pseudo-females using an identical pipeline as used by Halligan *et al.* (2013), as outlined above, using the same AFS prior.

Switch error rates in Shapeit2 are sensitive both to coverage and quality (per genotype and per variant) (Delaneau *et al.* 2013). We explored the effects of different filter parameters on the switch error rates produced by ShapeIt2 using the X-chromosomes of the pseudo-females. We filtered SNPs based on combinations of variant and genotype quality scores (QUAL and GQ, respectively) and on an individual’s sequencing depth (DP) (Table S1). For the individual-specific statistics (DP and GQ), if a single individual failed a particular filter, then that SNP was not included in further analyses. By comparing the known X-chromosome haplotypes and those inferred by ShapeIt2, we calculated switch error rates as the ratio of incorrectly resolved heterozygous SNPs to the total number of heterozygous SNPs for each pseudo-female individual. We used these results to choose filter parameters to apply to the autosomal data that generated a low switch error rate in ShapeIt2, while maintaining a high number of heterozygous SNPs. We obtained 20 phased haplotypes for each of the 19 mouse autosomes. With these, we estimated the recombination rate landscape for *M. m. castaneus.*

### Estimating recombination maps and validation of the approach

LDhelmet (v1.7;Chan *et al.* 2012) generates a sex-averaged map of recombination rates from a sample of haplotypes that are assumed to be drawn from a randomly mating population. Briefly, LDhelmet examines patterns of LD in a sample of phased chromosomal regions and uses a composite likelihood approach to infer recombination rates that are best supported between adjacent SNPs. LDhelmet appears to perform well for species with large effective population size (*N_e_*) and has been shown to be robust to the effects of selective sweeps, which may be prevalent and reduce diversity in and around functional elements of the *M. m. castaneus* genome (Halligan *et al.* 2013). However, the analyses conducted by Chan *et al.* (2012), in which the software was tested, were performed with a larger number of haplotypes than we have in our sample. To assess whether our smaller sample size gives reliable recombination maps, we validated and parameterized LDhelmet using simulated datasets.

A key parameter in LDhelmet is the block penalty, which determines the extent by which likelihood is penalized by spatial variation in the recombination rate, such that a high block penalty results in a smoother recombination map. We performed simulations to determine the block penalty that leads to the most accurate estimates of the recombination rate in chromosomes that have levels of diversity and base content similar to *M. m. castaneus.* Chromosomes with constant values of *ρ* (*4N_e_r*) ranging from 2 × 10^-6^ to 2 × 10^1^ were simulated in SLiM v1.8 (Messer 2013). For each value of *ρ*, 0.5Mbp of neutrally evolving sequence was simulated for populations of *N* = 1,000 diploid individuals. Mutation rates in the simulations were set using the compound parameter *θ* = *4N_e_ μ,* where *μ* is the per-base, per-generation mutation rate. The mutation and recombination rates of the simulations were scaled to *θ/4Ν* and *ρ/4N,* respectively. *θ* was set to 0.01 for all simulations, as this is close to the genome-wide average for our data, based on pairwise differences. Simulations were run for 10,000 generations to achieve equilibrium levels of polymorphism, at which time 10 diploid individuals were sampled from the population. Each simulation was repeated 20 times, resulting in 10Mbp of sequence for each value of *ρ*. The SLiM output files were converted to sequence data, suitable for analysis by LDhelmet, using a custom Python script that incorporated the mutation rate matrix estimated for non-CpG prone sites in *M. m. castaneus* (see below). We inferred recombination rates from the simulated data in windows of 4,400 SNPs with a 200 SNP overlap between windows, following (Chan *et al.* 2012). We analyzed the simulated data using LDhelmet with block penalties of 10, 25, 50 and 100. The default parameters of LDhelmet are tuned to analyze *Drosophila melanogaster* data (Chan *et al.* 2012). Since the *D. melanogaster* population studied by Chan *et al.* (2012) has comparable levels of genetic diversity to *M. m. castaneus* we used the defaults for all other parameters, other than the block penalty and estimate of *θ*.

Errors in phase inference, discussed above, may bias our estimates of the recombination rate, since they appear to break apart patterns of LD. We assessed the impact of these errors on recombination rate inference by incorporating them into the simulated data at a rate estimated from the pseudo-female individuals. For each of the 10 individuals drawn from the simulated populations, switch errors were randomly introduced at heterozygous positions at the rate estimated using the chosen SNP filter set (see Results). We then inferred the recombination rates, as above, for the simulated population using these error-prone data. We assessed the effect of switch errors on recombination rate inference by comparing estimates based on the simulated data both with and without switch errors. It is worth noting that there is the potential for switch errors to undo crossing-over events, reducing inferred recombination rates, if they affect heterozygous SNPs that are breakpoints of recombinant regions.

### Recombination rate estimation for M. m. castaneus

We used LDhelmet (Chan *et al.* 2012), to estimate recombination rates for each of the *M. m. castaneus* autosomes. It is well established that autosomal recombination rates differ between the sexes in *M musculus* (Cox *et al.* 2009; Liu *et al.* 2014). A drawback of LD-based approaches is that they give sex-averaged recombination rates.

We used both *M. famulus* and *R. norvegicus* as outgroups to assign ancestral alleles to polymorphic sites. LDhelmet incorporates both the mutation matrix and a prior probability on the ancestral allele at each variable position as parameters in the model. We obtained these parameters as follows. For non-CpG prone polymorphic sites, if the outgroups shared the same allele, we assigned that allele as ancestral and these sites were then used to populate the mutation matrix, following Chan *et al.* (2012). This approach ignores the possibility of both back mutation and homoplasy. To account for this uncertainty, LDhelmet incorporates a prior probability on the ancestral base. Following Singhal *et al.* (2015), at resolvable sites (i.e. when both outgroups agreed), the ancestral base was given a prior probability of 0.91, with 0.03 assigned to each of the three remaining bases. This was done to provide high confidence in the ancestral allele, but to also include the possibility of ancestral allele misinference. At unresolved sites (i.e., if the outgroup alleles did not agree or there were alignment gaps in either outgroup), we used the stationary distribution of allele frequencies from the mutation rate matrix as the prior (Table S2).

We analyzed a total of 43,366,235 SNPs in LDhelmet to construct recombination maps for each of the *M. m. castaneus* autosomes. Following Chan *et al.* (2012), windows of 4,400 SNPs, overlapping by 200 SNPs on either side, were analysed. We ran LDhelmet with a block penalty of 100 for a total of 1,000,000 iterations, discarding the first 100,000 as burn-in. The block penalty value was chosen to obtain a conservatively estimated recombination map, on the basis of the simulation analysis. We analyzed all sites that passed the filters chosen using the pseudo-female phasing regardless of CpG status; note that excluding CpG-prone sites removes ~50% of the available data and thus would substantially reduce the power to infer recombination rates. We assumed *θ* = 0.01, the approximate genome-wide level of neutral diversity in *M. m. castaneus*, and included ancestral allele priors and the mutation rate matrix for non-CpG sites as parameters in the model. Following the analyses, we removed overlapping SNPs and concatenated SNP windows to obtain recombination maps for whole chromosomes.

It is worthwhile noting that our map was constructed with genotype calls made using the mm9 version of the mouse reference genome. This version was released in 2007 and there have been subsequent versions released since then. However, previously published genetic maps for *M. musculus* were constructed using mm9, so we used that reference to make comparisons (see below).

### Comparison to previously published maps

The recombination rate map inferred for *M. m. castaneus* was compared with two previously published genetic maps for *M. musculus.* The first map was generated by analyzing the inheritance patterns of markers in crosses between inbred lines (Cox *et al.* 2009)(downloaded from http://cgd.jax.org/mousemapconverter/). Hereafter, this map shall be referred to as the Cox map. The second map was generated by Brunschwig *et al.* (2012), by analyzing SNPs in a large number of inbred mouse lines using LDhat (Auton and Mcvean 2007), the software upon which LDhelmet is based (available at http://www.genetics.org/content/early/2012/05/04/genetics.112.141036). Hereafter, this map shall be referred to as the Brunschwig map. Both maps were generated using classical strains of laboratory mice, which are predominantly of *M. m. domesticus* origin (Yang *et al.* 2011). Both the Brunschwig and Cox maps were constructed using far fewer markers than the present study, ~250,000 and ~10,000 SNPs, respectively.

Recombination rates in the Brunschwig map and our *castaneus* map were inferred in terms of the population recombination rate (*ρ* = *4N_e_r*), units that are not directly convertible to centimorgans (cM), but were converted to cM/Mb for comparison purposes using frequency weighted means, as follows. Both LDhat and LDhelmet give estimates of *ρ* (per Kbp and bp, respectively) between pairs of adjacent SNPs. To account for differences in the physical distance between adjacent SNPs when calculating cumulative *ρ*, we used the number of bases between a pair of SNPs to weight that pair’s contribution to the sum. By setting the total map distance for each chromosome to be equal to those found by Cox *et al.* (2009), we scaled the cumulative *ρ* at each analyzed SNP position to cM values.

At the level of whole chromosomes, we compared mean recombination rates from the *castaneus* map with several previously published maps. The frequency-weighted mean recombination rates (in terms of *ρ*) for each of the autosomes from the *castaneus* and Brunschwig maps were compared with the cM/Mb values obtained by Cox *et al.* (2009) as well as independent estimates of the per chromosome recombination rates from Jensen-Seaman *et al.* (2004). Pearson correlations were calculated for each comparison. Population structure in the inbred line data analyzed by Brunschwig *et al.* (2012) may have elevated LD, thus downwardly biasing estimates of *ρ*. To investigate this, we divided the frequency-weighted mean recombination rates per chromosome from the *castaneus* and Brunschwig maps by the rates given in Cox *et al.* (2009) to obtain estimates of effective population size.

At a finer scale, we compared variation in recombination rates across the autosomes in the different maps using windows. We calculated Pearson correlations between the frequency weighted-mean recombination rates (in cM/Mb) in non-overlapping windows for the *castaneus,* Cox and Brunschwig maps. The window size considered may affect the correlation between maps, so we calculate Pearson correlations in windows of 1Mbp to 20Mbp in size. For visual comparison of the *castaneus* and Cox maps, we plotted recombination rates in sliding windows of 10Mbp, offset by 1Mb.

### Examining the correlation between nucleotide diversity and recombination rate

There is evidence that natural selection is pervasive in the protein-coding genes and conserved non-coding elements in the murid genome (Halligan *et al.* 2010; Halligan *et al.* 2011; Halligan *et al.* 2013). Directional selection acting on selected sites within exons may reduce diversity at linked neutral sites through the processes of background selection and/or selective sweeps. These processes have the largest effect in regions of low recombination, and can therefore generate positive correlations between diversity and the recombination rate, as has been observed in multiple species (Cutter and Payseur 2013). We used our *castaneus* map to examine the relationship between nucleotide diversity and recombination rates as follows. We obtained the coordinates of the canonical spliceforms of protein coding genes, orthologous between mouse and rat from Ensembl Biomart (Ensembl Database 67; http://www.ensembl.org/info/website/archives/index.html). We calculated the frequency-weighted mean recombination rate and the GC content for each gene. Using the approximate *castaneus* reference, described above, and the outgroup alignment, we obtained the locations of 4-fold degenerate synonymous sites. If a site was annotated as 4-fold in all three species considered, it was used for further analysis. We removed poor quality alignments between mouse and rat, exhibiting a spurious excess of diverged sites, where >80% of sites were missing. We also excluded five genes that were diverged at all non-CpG prone 4-fold sites, as it is likely that these also represent incorrect alignments. After filtering, there were a total of 18,171 protein-coding genes for analysis.

We examined the correlation between local recombination rates in protein coding genes with nucleotide diversity and divergence. Variation in the mutation rate across the genome may influence genome-wide analyses of nucleotide polymorphism, so we also examined the correlation between the ratio of nucleotide diversity and divergence from *R. norvegicus* at neutral sites and the rate of recombination. We used non-parametric Kendall rank correlations for all comparisons.

All analyses were conducted using custom Python scripts, except correlation analyses which were conducted using R (R Core Team 2016).

## Results

### Phasing SNPs and estimating the switch error rate

In order to infer recombination rates from our sample of individuals, we required phased SNPs. Taking advantage of the high sequencing depth of the sample, we phased SNPs using ShapeIt2, an approach that makes use of both LD and sequencing reads to resolve haplotypes. We phased each of the mouse autosomes, giving a total of 43,366,235 SNPs for estimation of recombination rates.

By constructing pseudo-female individuals, we quantified the switch error rate incurred when inferring phase from our data. After filtering of variants, ShapeIt2 achieved low switch error rates for all parameter combinations tested (Table S1). We chose a set of filters (GQ > 15, QUAL > 30) that resulted in a mean switch error rates across the three pseudo-females of 0.46% (Table S1). More stringent filtering resulted in slightly lower mean switch error rates, but also resulted in the removal of many more variants from the dataset (Table S1), thus reducing power to resolve recombination rates in downstream analyses.

### Simulations to validate LDhelmet for the population sample ofM. m. castaneus

We assessed the performance of LDhelmet when applied to our dataset by simulation. In the absence of switch errors, LDhelmet accurately infers the average recombination rate down to values of *ρ*/bp = 2×10^-4^ (Figure 1). Below this value, LDhelmet overestimated the scaled recombination rate for the simulated populations (Figure 1). With switch errors incorporated into simulated data, LDhelmet accurately estimated *ρ*/bp in the range 2×10^-3^ to 2×10^2^. When the true *ρ*/bp was <2×10^-3^, however, LDhelmet overestimated the mean recombination rate for 0.5Mbp regions (Figure 1). This behavior was consistent for all block penalties tested (Figure S1). Given that the simulations incorporated the mutation rate matrix (Table S2) and mutation rate (*θ* = *4Ν_e_ μ*) estimated for *M. m. castaneus* we concluded that LDhelmet is applicable to the dataset of 10 *M. m. castaneus* individuals sequenced by Halligan *et al.* (2013).

**Figure 1.**
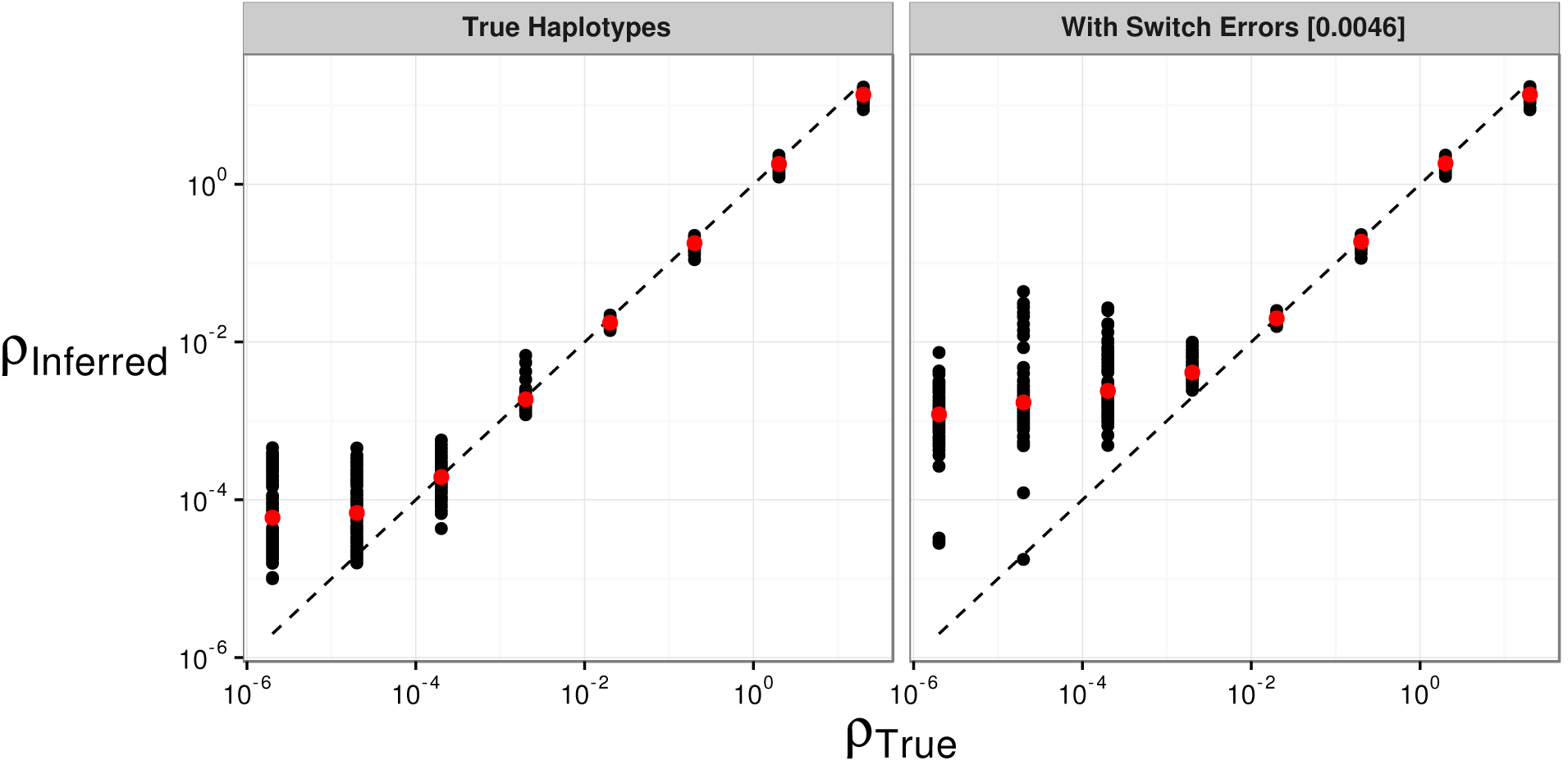
The effect of switch errors on the mean recombination rate inferred using LDhelmet with a block penalty of 100. Each black point represents results for a window of 4,000 SNPs, with 200 SNPs overlapping between adjacent windows, using sequences simulated in SLiM for a constant value of ρ/bp. Red points are mean values. Switch errors were randomly incorporated at heterozygous SNPs with probability 0.0046. The dotted line shows the value for inferred=true.

### Recombination rates across the M. m. castaneus autosomes

A recombination rate map for each *M. m. castaneus* autosome was constructed using LDhelmet. We analyzed a total of 43,366,235 phased SNPs across the 19 mouse autosomes. The frequency weighted mean value of *ρ*/bp for all autosomes was 0.009. This value is greater than the lower detection limit suggested by both the simulations with and without switch errors (Figure 1).

We assessed variation in whole-chromosome recombination rates between our LD-based *castaneus* map and direct estimates of recombination rates published in earlier studies. Comparing the mean recombination rates for whole chromosomes provides us with a baseline comparison for which we have an *a priori* expectation: we expect that chromosome 19, the shortest in physical length, should have the highest mean recombination rate, since at least one crossing over event is required per meiosis per chromosome in mice. This has been demonstrated in previous studies of recombination in *M.* musculus (Jensen-Seaman *et al.* 2004; Cox *et al.* 2009). Indeed, we find that the frequency-weighted mean recombination rate for chromosome 19 is the highest among the autosomes (Table 1). We also found that the frequency-weighted mean recombination rates for each of the autosomes were highly correlated with the direct estimates given in Jensen-Seaman *et al.* (2004) (Pearson correlation = 0.66, *p* = 0.002) and Cox *et al.* (2009) (Pearson correlation = 0.88, *p* < 0.0001), suggesting that our analysis captures real variation in recombination rates at the scale of whole chromosomes in the *M. m. castaneus* genome.

**Table 1.** Summary of sex-averaged recombination rates estimated for the *M. m castaneus* autosomes compared with the rates from Brunschwig *et al.* (2012) and Cox *et al.* (2009). Rates for the *castaneus* and Brunschwig maps are presented in terms of 4*N_e_r*/bp. Estimates of *N_e_* were obtained by assuming the recombination rates from Cox *et al.* (2009).

| Chromosome | Cox<br>cM/Mb | *castaneus* |  | Brunschwig |  |
| --- | --- | --- | --- | --- | --- |
| | | Freq.<br>Weighted<br>Mean | $N_e$ Estimate | Freq.<br>Weighted<br>Mean | $N_e$ Estimate |
| 1 | 0.50 | 0.0079 | 395,000 | 0.000015 | 745 |
| 2 | 0.57 | 0.0088 | 386,000 | 0.000015 | 653 |
| 3 | 0.52 | 0.0083 | 400,000 | 0.000014 | 693 |
| 4 | 0.56 | 0.0091 | 408,000 | 0.000020 | 889 |
| 5 | 0.59 | 0.0090 | 382,000 | 0.000015 | 646 |
| 6 | 0.53 | 0.0089 | 421,000 | 0.000015 | 728 |
| 7 | 0.58 | 0.0100 | 429,000 | 0.000019 | 801 |
| 8 | 0.58 | 0.0094 | 404,000 | 0.000014 | 610 |
| 9 | 0.61 | 0.0096 | 394,000 | 0.000018 | 749 |
| 10 | 0.61 | 0.0096 | 392,000 | 0.000023 | 928 |
| 11 | 0.70 | 0.0102 | 365,000 | 0.000019 | 689 |
| 12 | 0.53 | 0.0089 | 420,000 | 0.000019 | 897 |
| 13 | 0.56 | 0.0095 | 426,000 | 0.000014 | 629 |
| 14 | 0.53 | 0.0084 | 395,000 | 0.000013 | 632 |
| 15 | 0.56 | 0.0083 | 371,000 | 0.000024 | 1,080 |
| 16 | 0.59 | 0.0091 | 386,000 | 0.000017 | 721 |
| 17 | 0.65 | 0.0087 | 335,000 | 0.000052 | 2,020 |
| 18 | 0.66 | 0.0098 | 371,000 | 0.000021 | 785 |
| 19 | 0.94 | 0.0122 | 323,000 | 0.000026 | 681 |
| Mean |  | 0.0092 |  | 0.000020 |  |

### Comparison of the M. m. castaneus map to maps constructed using inbred lines

We compared the intra-chromosomal variation in recombination rates between our *castaneus* map and previously published maps. Figure 2 shows the variation in recombination rates across the largest and smallest autosomes in the mouse genome, chromosomes 1 and 19, respectively. It is clear that the *castaneus* and Cox maps are very similar (see also Figure S2 showing a comparison of all autosomes). Correlation coefficients between the maps are >0.8 for window sizes of 8Mbp and above (Figure 3). Although the overall correlation between the *castaneus* and Cox maps is high (Figure 3), there were several regions of the genome that substantially differ, for example in the centre of chromosome 9 (Figure S2). The Cox and *castaneus* maps are more similar to one another than either are to the Brunschwig map (Figure 3), presumably because the Brunschwig map was constructed with a sample of 60 inbred mouse strains. Population structure in the lines or the subspecies from which they were derived would elevate LD, resulting in downwardly-biased chromosome-wide values of *ρ*. This is also reflected in the *N_e_* values estimated from the frequency-weighted average recombination rates for each chromosome. The estimates of *N_e_* are substantially different between the *castaneus* and Brunschwig maps, i.e. the *castaneus* estimates are consistently ~500x higher (Table 1). The estimates of *N_e_* from the *castaneus* map are in broad agreement with the estimates of *N_e_* based on polymorphism data (Geraldes *et al.* 2008).

**Figure 2.**
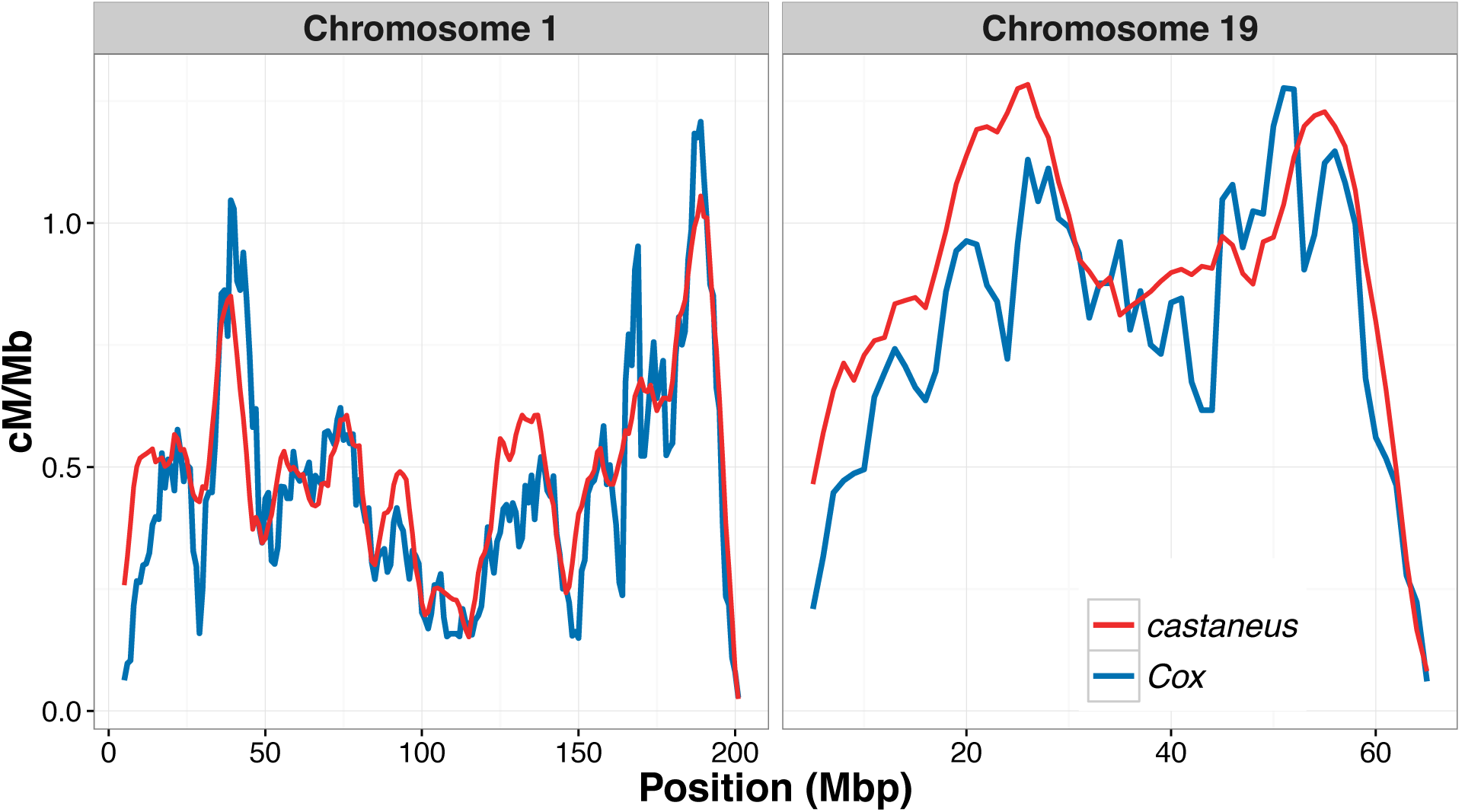
Comparison of the sex-averaged recombination rate inferred for chromosomes 1 and 19 of *M. musculus castaneus* using LDhelmet in red and those estimated from the pedigree-based study of Cox *et al.* (2009) in blue. Recombination rates in units of cM/Mb for the *castaneus* map were obtained by setting the total genetic lengths for each chromosome to the corresponding lengths from Cox *et al.* (2009).

**Figure 3.**
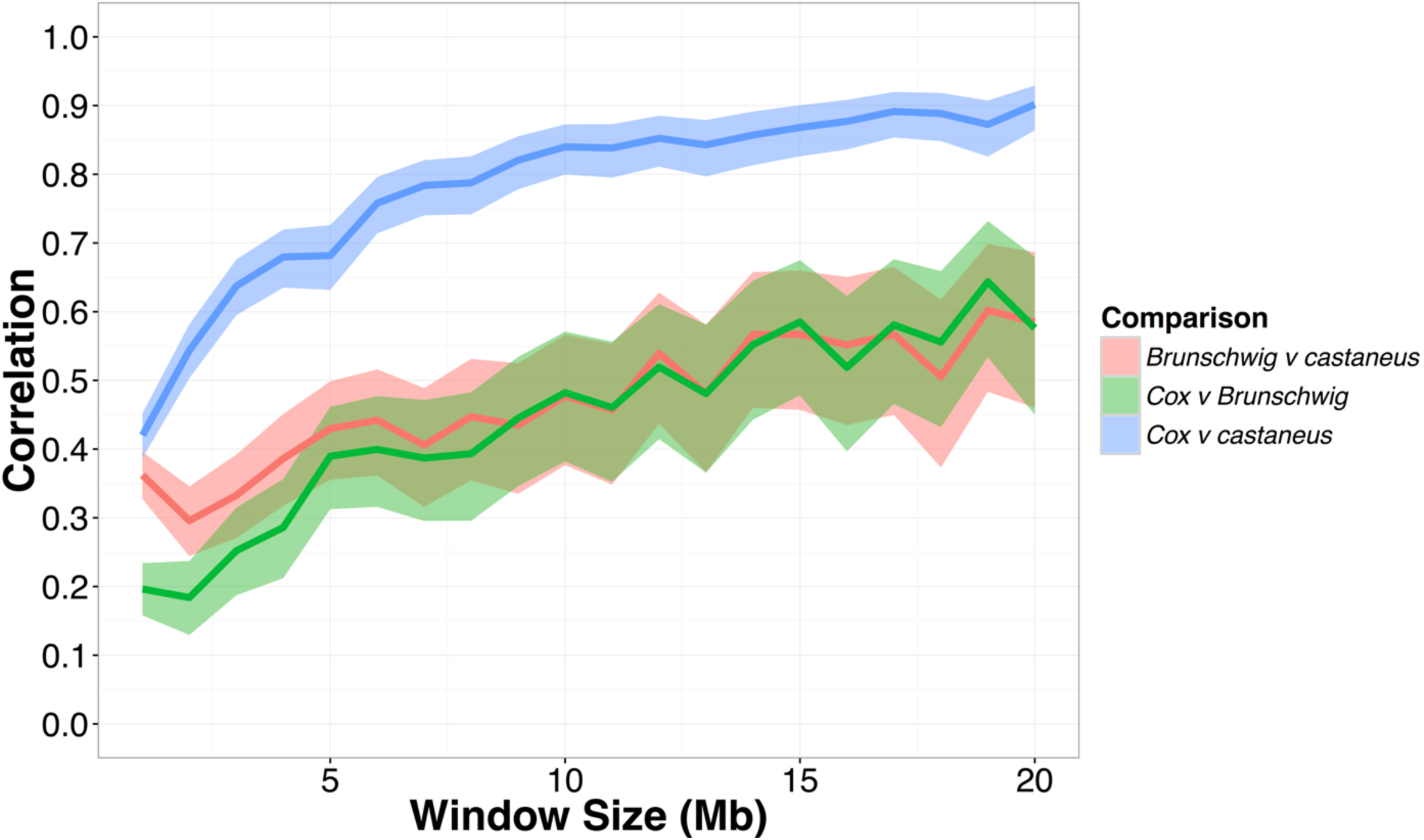
Pearson correlation coefficients between the recombination map inferred for *M. m. castaneus,* the Brunschwig *et al.* (2012) map and the Cox *et al.* (2009) map. Correlations were calculated in non-overlapping windows of varying size across all autosomes. Confidence intervals (95%) are indicated by shading around each line.

### Correlations between recombination rate and properties of protein coding genes in M. m. castaneus

By examining the correlation between genetic diversity and recombination rate, we determined whether our map captures variation in *N_e_* across the genome. We found that recombination rates at protein coding genes are significantly and positively correlated with levels of neutral genetic diversity (Table 2), at all sites regardless of base context and at non-CpG-prone sites only (Table 2). Divergence from the rat at 4-fold sites was also significantly and positively correlated with recombination rate when analyzing all sites. However, for non-CpG-prone sites we found a small negative correlation (Table 2). There was also a significant and positive relationship between recombination rate and a gene’s GC content (*τ* = 0.125, *p* < 2.2×10^-16^). The correlation between recombination rate and neutral diversity divided by divergence from the rat was both positive and significant, regardless of base context (Table 2; Figure S3). This indicates that natural selection may have a role in reducing diversity via hitchhiking and/or background selection.

**Table 2.** Correlation coefficients between recombination rate and pairwise nucleotide diversity and divergence from the rat at 4-fold degenerate sites for protein coding genes. Non-parametric Kendall correlations were calculated for non-CpG prone sites and for all sites, regardless of base context. All coefficients shown are highly significant (*p* < 10^-10^).

|  | Non-CpG | All Sites |
| --- | --- | --- |
| Nucleotide diversity ($\pi$) | 0.09 | 0.20 |
| Divergence to rat ($d_{rat}$) | -0.04 | 0.06 |
| Corrected diversity ($\pi/d_{rat}$) | 0.10 | 0.18 |

## Discussion

By constructing fine-scale maps of the recombination rate for the autosomes of *M. m. castaneus,* we have shown that there is a high degree of similarity between the recombination landscape for wild-caught mice and their laboratory counterparts. Our map captures variation in the recombination rate, similar to that observed in a more traditional linkage map, at the level of both whole chromosomes and genomic windows of varying size.

Recombination landscapes inferred using coalescent approaches, as in this study, reflect ancestral variation in recombination rates. We show that this ancestral variation is highly correlated with contemporaneous recombination rates in inbred mice of a different subspecies, suggesting that the broad-scale variation in recombination rate has not evolved dramatically since the subspecies diverged around 500,000 years ago (Geraldes *et al.* 2008). At a finer scale, however, Smagulova *et al.* (2016) showed that there is considerable variation in the locations of recombination hotspots between the *M. musculus* subspecies. Their findings, taken together with ours, parallel results in hominids and the great-apes, which suggest that, although the locations of recombination hotspots are strongly diverged between species, broad-scale patterns of recombination rate are relatively conserved (Lesecque *et al.* 2014; Stevison *et al.* 2015). However, there do seem to be multiple regions of the genome that distinguish *M. m. castaneus* and *M. m. domesticus.* For example, we observe peaks in recombination rate for *M. m. castaneus* on chromosomes 4, 5, 14 and 15 that are not present in the Cox map (Figure S2). These results are seemingly consistent with those of Dumont *et al.* (2011), who found that there are significant differences in genetic length between *M. m. castaneus* and *M. m. musculus* (when crossed to *M. m. domesticus*) in multiple regions of the genome (though a large proportion of the differences they detected were on the X-chromosome, which was not analyzed in our study). Performing a comparative analysis of recombination rates in the different subspecies of house mice, as well as sister species, using LD-based methods would help elucidate the time-scale of recombination rate evolution in wild mice.

The *castaneus* map constructed in this study appears to be more similar to the Cox map than the Brunschwig map (Figure 3). There are number of potential reasons for this. Firstly, we used a much larger number of markers to resolve recombination rates than Brunschwig *et al.* (2012), giving us more power to capture variation in the recombination rate. Secondly, it seems probable that population structure within and between the inbred and wild-derived lines studied by Brunschwig *et al.* (2012) could have resulted in biased estimates of the recombination rate. By dividing the mean estimated *ρ*/bp values (inferred using LDhelmet) for each chromosome by the corresponding recombination rate estimated from crosses (Cox *et al.* 2009), we showed that *N_e_* estimates from the Brunschwig map are much lower than estimates based on our map (Table 1). This is consistent with the presence of elevated LD between the SNPs in the inbred lines analyzed by Brunschwig *et al.* (2012). It should be noted, however, that the estimates of *N_e_* will be biased, as *θ = 4Ν_e_ μ* is a parameter in both LDhat and LDhelmet. In spite of this potential bias, the differences in *N_e_* estimated from the Brunschwig and *castaneus* maps shown in Table 1 are striking, given that the ancestral effective population sizes of *M. m. domesticus* and *M. m. castaneus* are expected to be ~150,000 and ~350,000, respectively (Geraldes *et al.* 2008). The Brunschwig map does, however, capture true variation in recombination rates, because their map is also highly correlated with the Cox map (Pearson correlation >0.6) for all genomic windows wider than 8Mbp (Figure 3). Indeed, Brunschwig *et al.* (2012) showed by simulation that hotspots are detectable by analysis of inbred lines and validated their inferred hotspots against the locations of those observed in crosses among classical strains of *M. m. domesticus* (Smagulova *et al.* 2011). This suggests, that while estimates of the recombination rate in the Brunschwig *et al.* (2012) map may have been downwardly biased by population structure, variation in the rate and locations of hotspots were still accurately detected in their study.

We obtained an estimate of the switch error rate, taking advantage of the hemizygous sex chromosomes of males present in our sample. This allowed us to assess the extent by which switch errors affected our ability to infer recombination rates in *M. m. castaneus.* It should be noted, however, that our inferred switch error rate may not fully represent that of the autosomes. This is because multiple factors influence the ability to phase variants using ShapeIt2 (i.e. LD, SNP density, sample size, depth of coverage and read length) and some of these factors differ between the X-chromosome and the autosomes. Firstly, as the sex-averaged recombination rate for the X-chromosome is expected to be 3/4 that of the autosomes, it likely has elevated LD, and thus there will be higher power to infer phase. In contrast, the level of X-linked nucleotide diversity in *M. m. castaneus* is approximately one half that of the autosomes (Kousathanas *et al.* 2014), and thus there would be a higher probability of phase informative reads on the autosomes. While it is difficult to assess whether the switch error rates we estimated from the X-chromosome analysis will be the same as on the autosomes, the analysis allowed us to explore the effects of different SNP filters on the error rate.

By simulating the effect of switch errors on estimates of the recombination rate, we inferred the range over which *ρ*/bp is accurately estimated in our data. Switch errors appear identical to legitimate crossing-over events and, if they are randomly distributed along chromosomes, a specific rate of error will resemble a constant rate of crossing over. The rate of switch error will then determine a detection threshold below which recombination cannot be accurately inferred. We introduced switch errors at random into the simulation data and estimates of *ρ*/bp obtained from these datasets reflect this detection threshold; below 2×10^-3^ *ρ*/bp, we found that LDhelmet consistently overestimates the recombination rate in the presence of switch errors (Figure 1; Figure S1). This highlights a possible source of bias affecting LD-based recombination mapping studies using inferred haplotypes. In a recent study, Singhal *et al.* (2015) showed that the power to detect recombination hotspots is reduced when the recombination rate in the regions surrounding a hotspot is low. Though we did not attempt to locate recombination hotspots in this study, our findings and those of Singhal *et al.* (2015) both suggest that error in phase inference needs to be carefully considered before attempting to estimate recombination rates and/or recombination hotspots using LD-based approaches.

Consistent with studies in a variety of organisms, we found a positive correlation between genetic diversity at putatively neutral sites and the rate of recombination. Both unscaled nucleotide diversity and diversity divided by divergence between mouse and rat, a proxy for the mutation rate, are positively correlated with recombination (Table 2). Cai *et al.* (2009) found evidence suggesting that recombination may be mutagenic, though insufficient to account for the correlations they observed between recombination and diversity. The Kendall correlation between *π/d_rat_* and recombination rate of 0.20 for all 4-fold sites, a value that is similar in magnitude to the corresponding value of 0.09 reported by Cai *et al.* (2009) in humans. The correlations we report may be downwardly biased, however, because switch errors may result in inflated recombination rates inferred for regions of the genome where the true recombination rate is low (see above). Genes that have recombination rates lower than the detection limit set by the switch error rate may be reported as having inflated *ρ*/bp (Figure 1; Figure S1), and this would have the effect of reducing correlation statistics. It is difficult to assess the extent of this bias, however, and in any case the correlations we observed between diversity and recombination suggest that our recombination map does indeed capture real variation in *N_e_* across the genome. This indicates that a recombination mediated process influences levels of genetic diversity. Previously, Halligan *et al.* (2013) showed that there are troughs in nucleotide diversity surrounding protein coding exons in *M. m. castaneus,* characteristic of natural selection acting within exons reducing diversity at linked sites. Their results and ours suggest pervasive natural selection in the genome of *M. m. castaneus.*

In conclusion, we find that sex-averaged estimates of the ancestral recombination landscape for *M. m. castaneus* are highly correlated with contemporary estimates of the recombination rate estimated from crosses of *M. m. domesticus* (Cox *et al.* 2009). It has been demonstrated previously that the turnover of hotspots has led to rapid evolution of fine-scale rates of recombination in the *M. musculus* subspecies complex (Smagulova *et al.* 2016). On a broad scale, however, our results suggest that the recombination landscape is very strongly conserved between the subspecies. In addition, our estimate of the switch-error rate implies that phasing errors leads to upwardly biased estimates of the recombination rate when the true recombination rate is low. This is a source of bias that should be assessed in future studies. Finally, we showed that the variation in recombination rate is positively correlated with genetic diversity, suggesting that natural selection reduces diversity at linked sites across the *M. m. castaneus* genome, consistent with the findings of Halligan et al (2013).

To further our understanding of the evolution of the rate of recombination in the house mouse we need to directly compare subspecies. The comparison of our results and previously published maps indicates that there is broad-scale agreement in recombination rates between *M. m. castaneus* and *M. m. domesticus.* In this study, we have assumed that inbred lines derived from *M. m. domesticus* reflect natural variation in recombination rates in that sub-species, though this is not necessarily the case. Population samples like the one studied here could be used to more clearly elucidate the recombination rate maps specific to the different subspecies. A broad survey of this kind would most efficiently be generated using LD-based approaches.

## Acknowledgements

We are grateful to Bettina Harr, Dan Halligan, Ben Jackson and Rory Craig for discussions and helpful comments on the manuscript. Tom Booker is supported by a BBSRC EASTBIO studentship. This project has received funding from the European Research Council (ERC) under the European Union’s Horizon 2020 research and innovation programme (grant agreement No. 694212). Rob Ness was funded by the BBSRC (BB/L00237X/1).

---

**Figure S1.**
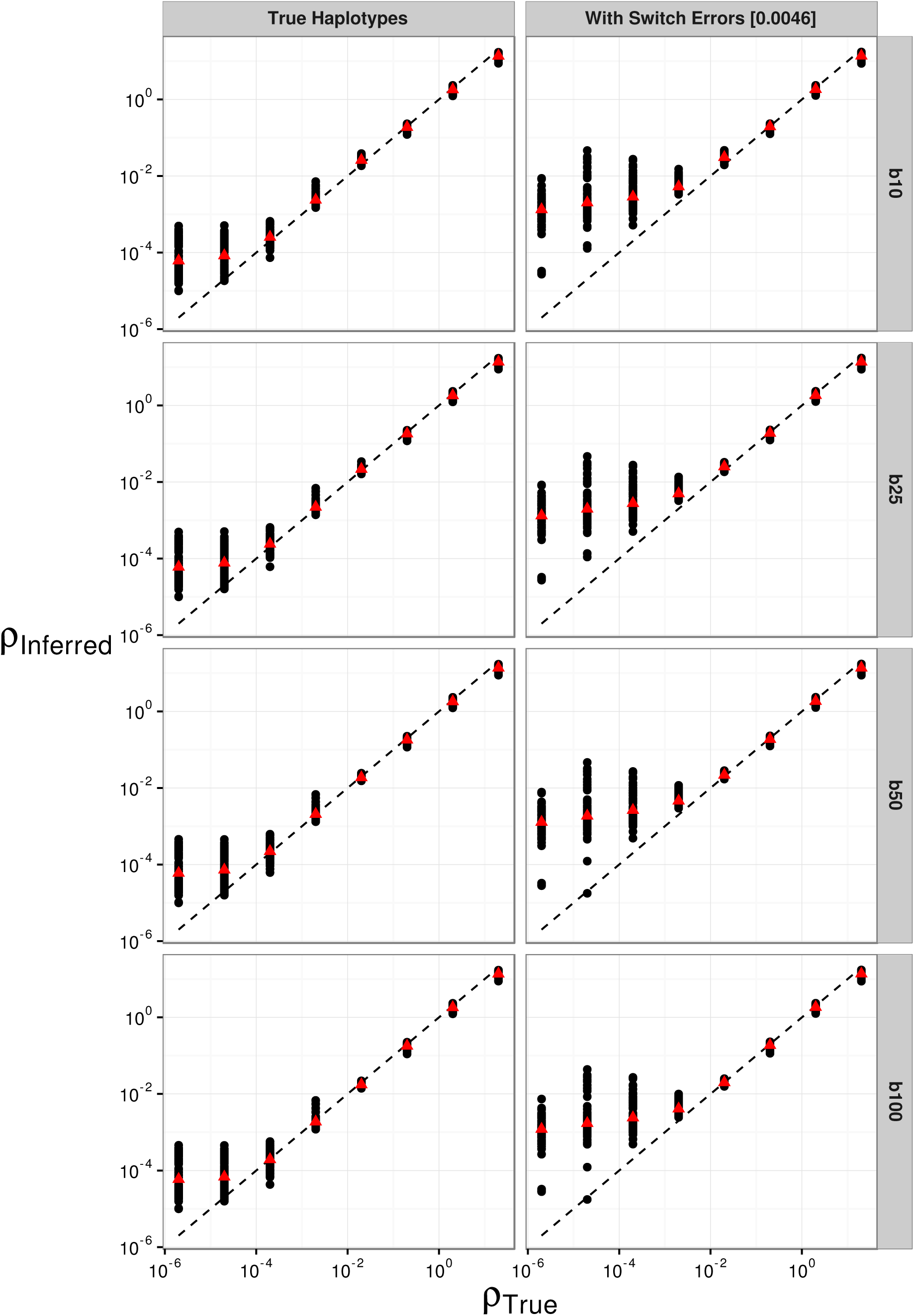
The effect of switch errors and block penalty on the mean recombination rate inferred using LDhelmet. Block penalties (b) of 10, 25, 50 and 100 were used, shown in the vertically ordered facets from top to bottom.

**Figure S2.**
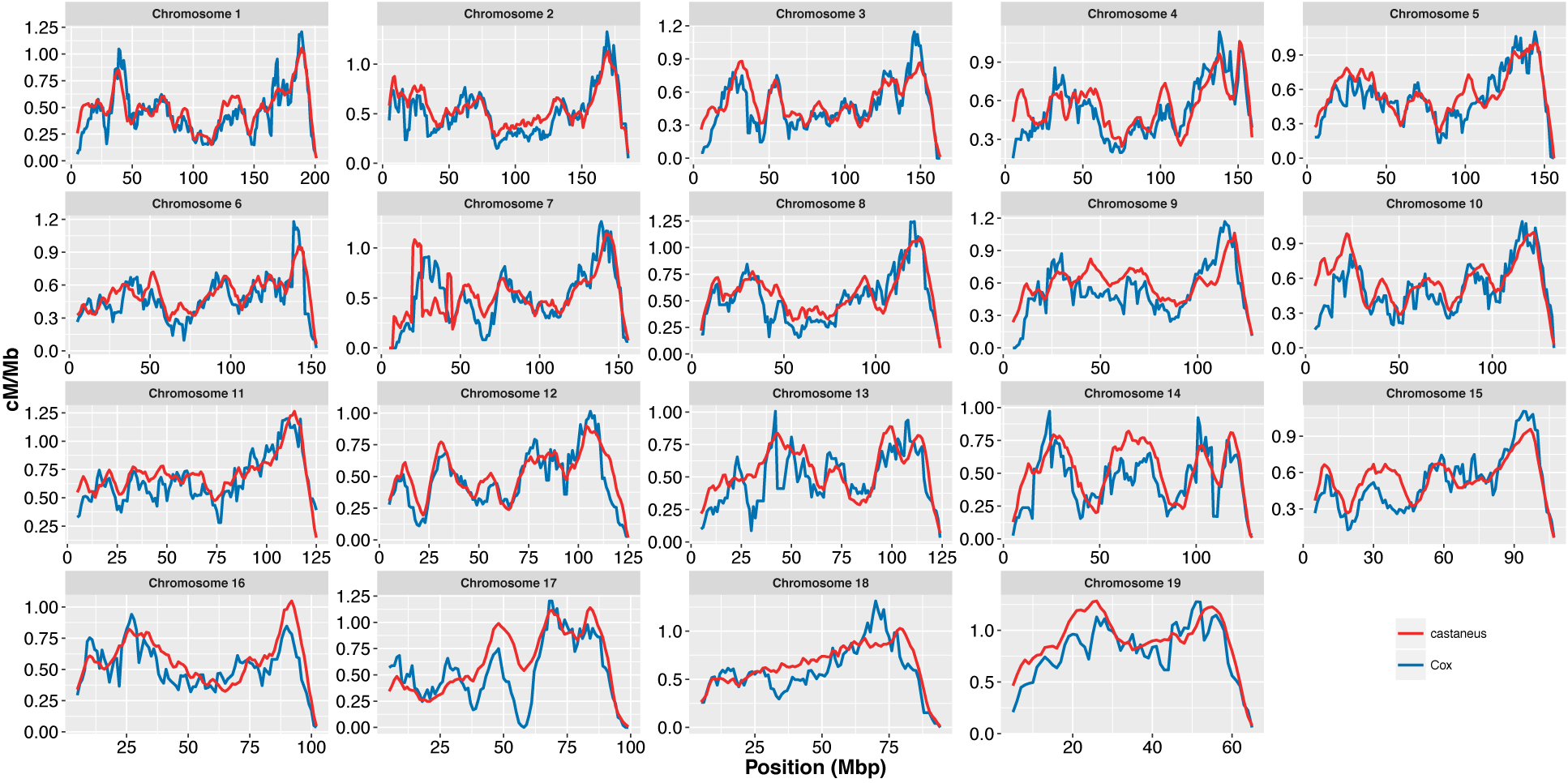
Comparison of recombination rates inferred for *M. m. castaneus* using LDhelmet and recombination rates reported by Cox et al (2009). Recombination rates in units of p/bp for the *castaneus* map were converted to cM/Mb by scaling using the genetic length of the corresponding chromosome in the Cox map.

**Figure S3.**
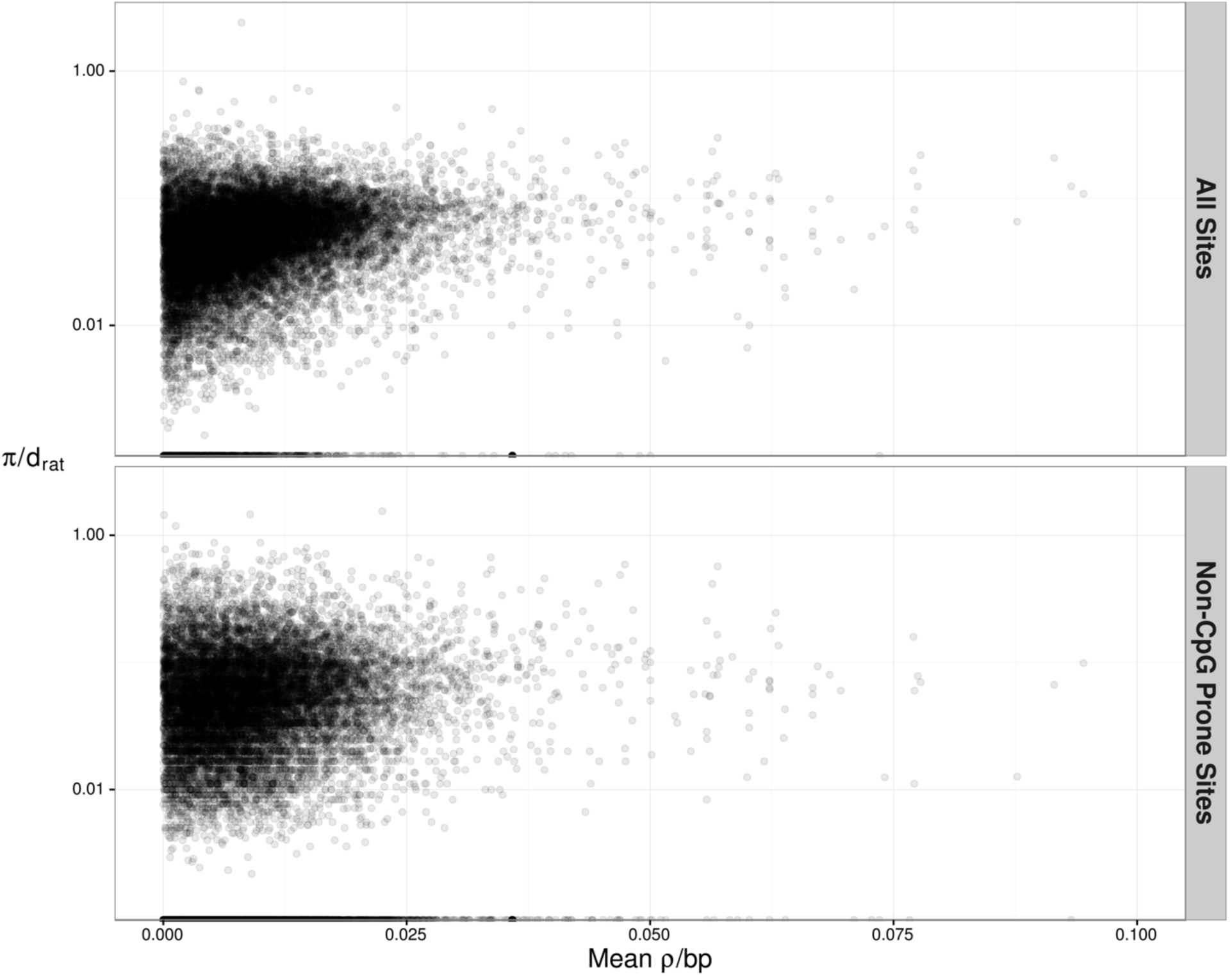
Nucleotide diversity divided by divergence from rat (*π/d_rat_*) *at* 4-fold degenerate synonymous sites plotted against the frequency weighted mean recombination rates (*ρ*/bp) for protein coding genes in *M. m. castaneus.* Correlation statistics are shown in the main text. For the purposes of visualization, the range of *ρ*/bp is restricted from 0 to 0.1. There are 19 genes that have mean p/bp >0.1, however.

**Table S1.** The effect of different filters on the frequency of switch errors in the haplotypes inferred based on the three pseudo-females. The values in the switch errors column are the raw numbers of switch errors and the total number of heterozygous SNPs on the X-chromsome. Variants with Quality (QUAL) <30 were excluded for all filter sets.

| Filter Set | HWE<sup>*</sup> | Min DP<sup>†</sup> | Max DP | Min GQ<sup>‡</sup> | Switch Errors |  |  | Switch Error Rate |
| --- | --- | --- | --- | --- | --- | --- | --- | --- |
|  |  |  |  |  | H40 | H46 | H62 |  |
| 1 | - | - | - | - | 5148 / 409486 | 4819 / 407422 | 5020 / 394778 | 0.0124 |
| 2 | <0.0002 | 10 | - | 15 | 1690 / 338592 | 1451 / 334111 | 1452 / 324199 | 0.0046 |
| 3 | <0.0002 | 10 | 100 | 5 | 2460 / 341744 | 2066 / 339508 | 2536 / 328998 | 0.0070 |
| 4 | <0.0002 | - | - | 40 | 523 / 288471 | 444 / 286636 | 550 / 281266 | 0.0018 |

<sup>\*</sup> HWE refers to the p-value for the Samtools Hardy-Weinberg equilibrium test below which variants were excluded.
<sup>†</sup> Depth of coverage per individual.
<sup>‡</sup> Per individual genotype quality scores.

**Table S2.** The normalized mutation rate matrix and stationary distribution of base frequencies estimated with two out-groups, *M. famulus* and *R. norvegicus,* using the method described by Chan *et al.* (2012).

|  | A | C | G | T |
| --- | --- | --- | --- | --- |
| A | 0.48 | 0.09 | 0.32 | 0.11 |
| C | 0.19 | 0.00 | 0.12 | 0.69 |
| G | 0.69 | 0.12 | 0.00 | 0.19 |
| T | 0.11 | 0.32 | 0.08 | 0.48 |
| ----- |  |  |  |  |
| Stationary<br>Distribution | 0.34 | 0.16 | 0.16 | 0.34 |

## Literature Cited

Auton, A. and G. McVean 2007 Recombination rate estimation in the presence of hotspots. Genome Res 17: 1219–1227.

Baines, J.F., and B. Harr, 2007 Reduced x-linked diversity in derived populations of house mice. Genetics 175: 1911–1921.

Baudat, F., J. Buard, C. Grey, A. Fledel-Alon, C. Ober et al., 2010 Prdm9 is a major determinant of meiotic recombination hotspots in humans and mice. Science 327: 836–840.

Baudat, F., Y. Imai and B. de Massy, 2013 Meiotic recombination in mammals: Localization and regulation. Nat Rev Genet 14: 794–806.

Brick, K., F. Smagulova, P. Khil, R.D. Camerini-Otero and G.V. Petukhova, 2012 Genetic recombination is directed away from functional genomic elements in mice. Nature 485: 642–645.

Brunschwig, H., L. Liat, E. Ben-David, R.W. Williams, B. Yakir et al., 2012 Fine-scale maps of recombination rates and hotspots in the mouse genome. Genetics 191: 757–764.

Cai, J.J., J. M. Macpherson, G. Sella and D.A. Petrov 2009 Pervasive hitchhiking at coding and regulatory sites in humans. PLoS Genet 5: e1000336.

Chan, A.H., P. A. Jenkins and Y.S. Song 2012 Genome-wide fine-scale recombination rate variation in drosophila melanogaster. PLoS Genet 8: e1003090.

Cox, A., C. L. Ackert-Bicknell, B.L. Dumont, Y. Ding, J.T. Bell et al., 2009 A new standard genetic map for the laboratory mouse. Genetics 182: 1335–1344.

Cutter, A.D., and B.A. Payseur 2013 Genomic signatures of selection at linked sites: Unifying the disparity among species. Nat Rev Genet 14: 262–274.

Davies, B., E. Hatton, N. Altemose, J.G Hussin, F. Pratto et al., 2016 Re-engineering the zinc fingers of prdm9 reverses hybrid sterility in mice. Nature 530: 171–176.

Delaneau, O., B. Howie, A.J. Cox, J.F. Zagury and J. Marchini 2013 Haplotype estimation using sequencing reads. Am J Hum Genet 93: 687–696.

Dumont, B.L., M.A. White, B. Steffy, T. Wiltshire and B.A. Payseur 2011 Extensive recombination rate variation in the house mouse species complex inferred from genetic linkage maps. Genome Res 21: 114–125.

Geraldes, A., P. Basset, B. Gibson, K.L. Smith, B. Harr et al., 2008 Inferring the history of speciation in house mice from autosomal, x-linked, y-linked and mitochondrial genes. Mol Ecol 17: 5349–5363.

Grey, C., P. Barthes, G. Chauveau-Le Friec, F. Langa, F. Baudat et al., 2011 Mouse prdm9 DNA-binding specificity determines sites of histone h3 lysine 4 trimethylation for initiation of meiotic recombination. PLoS Biol 9: e1001176.

Halligan, D.L., A. Kousathanas, R.W. Ness, B. Harr, L. Eory et al., 2013 Contributions of protein-coding and regulatory change to adaptive molecular evolution in murid rodents. PLoS Genet 9: e1003995.

Halligan, D.L., F. Oliver, A. Eyre-Walker, B. Harr and P.D. Keightley 2010 Evidence for pervasive adaptive protein evolution in wild mice. PLoS Genet. 6: e1000825.

Halligan, D. L., F. Oliver, J. Guthrie, K.C. Stemshorn, B. Harr et al., 2011 Positiveand negative selection in murine ultraconserved noncoding elements. Mol Biol Evol. 28: 2651–2660.

Hudson, R.R., 2001 Two-locus sampling distributions and their applications. Genetics. 159: 12.

Jensen-Seaman, M.I., T. S. Furey, B.A. Payseur, Y. Lu, K.M. Roskin et al., 2004 Comparative recombination rates in the rat, mouse and human genomes. Genome Res. 14: 528–538.

Johnston, S.E., C. Berenos, J. Slate and J.M. Pemberton 2016 Conserved genetic architecture underlying individual recombination rate variation in a wild population of soay sheep (ovis aries). Genetics. 203: 583–598.

Kousathanas, A., D. L. Halligan and P.D. Keightley 2014 Faster-x adaptive protein evolution in house mice. Genetics. 196: 1131–1143.

Lesecque, Y., S. Glemin, N. Lartillot, D. Mouchiroud and L. Duret, 2014 The red queen model of recombination hotspots evolution in the light of archaic and modern human genomes. PLoS Genet. 10: e1004790.

Li, H., and R. Durbin 2009 Fast and accurate short read alignment with burrows-wheeler transform. Bioinformatics. 25: 1754–1760.

Li, H., B. Handsaker, A. Wysoker, T. Fennell, J. Ruan et al., 2009 The sequence alignment/map format and samtools. Bioinformatics. 25: 2078–2079.

Liu, E.Y., A. P. Morgan, E.J. Chesler, W. Wang, G.A. Churchill et al., 2014 High-resolution sex-specific linkage maps of the mouse reveal polarized distribution of crossovers in male germline. Genetics. 197: 91–106.

McVean, G., P. Awadalla and P. Fearnhead 2002 A coalescent-based method for detecting and estimating recombination from gene sequences. Genetics. 160: 1231–1241.

McVean, G., S.R. Myers, S. Hunt, P. Deloukas, D.R. Bentley et al., 2004 The fine-scale structure of recombination rate variation in the human genome. Science. 304.

Messer, P.W., 2013 Slim: Simulating evolution with selection and linkage. Genetics. 194: 1037–1039.

Myers, S.R., R. Bowden, A. Tumian, R.E. Bontrop, C. Freeman et al., 2010 Drive against hotspot motifs in primates implicates the prdm9 gene in meiotic recombination. Science. 327.

Paigen, K., and P. Petkov 2010 Mammalian recombination hot spots: Properties, control and evolution. Nat Rev Genet. 11: 221–233.

Paigen, K., J. P. Szatkiewicz, K. Sawyer, N. Leahy, E.D. Parvanov et al., 2008 The recombinational anatomy of a mouse chromosome. PLoS Genet. 4: e1000119.

R Core Team, 2016 R: A language and environment for statistical computing., pp. R Foundation for Statistical Computing Vienna Austria.

Schwartz, J.J., D. J. Roach, J.H. Thomas and J. Shendure, 2014 Primate evolution of the recombination regulator prdm9. Nat Commun. 5: 4370.

Singhal, S., E. Leffler, K. Sannareddy, I. Turner, O. Venn et al., 2015 Stable recombination hotspots in birds. Science. 350: 6.

Smagulova, F., K. Brick, P. Yongmei, R.D. Camerini-Otero and G.V. Petukhova 2016 The evolutionary turnover of recombiantion hotspots contributes to speciation in mice. Genes & Development. 30: 277–280.

Smagulova, F., I. V. Gregoretti, K. Brick, P. Khil, R.D. Camerini-Otero et al., 2011 Genome-wide analysis reveals novel molecular features of mouse recombination hotspots. Nature. 472: 375–378.

Smukowski, C.S., and M.A. Noor, 2011 Recombination rate variation in closely related species. Heredity (Edinb). 107: 496–508.

Smukowski Heil, C.S., C. Ellison, M. Dubin and M.A. Noor, 2015 Recombining without hotspots: A comprehensive evolutionary portrait of recombination in two closely related species of drosophila. Genome Biol Evol. 7: 2829–2842.

Stevison, L.S., K. B. Hoehn and M.A. Noor, 2011 Effects of inversions on within-and between-species recombination and divergence. Genome Biol Evol. 3: 830–841.

Stevison, L.S., A. E. Woerner, J.M. Kidd, J.L. Kelley, K.R. Veeramah et al., 2015 The time scale of recombination rate evolution in great apes. Mol Biol Evol.

Wang, R.J., M. M. Gray, M.D. Parmenter, K.W. Broman and B.A. Payseur 2017 Recombination rate variation in mice from an isolated island. Mol Ecol. 26: 457–470.

Winckler, W., S.R. Myers, D.J. Richter, R.C. Onofrio, G.J. McDonald et al., 2005 Comparison of fine-scale recombination rates in humans and chimpanzees. Science. 308.

Yang, H., J. R. Wang, J.P. Didion, R.J. Buus, T.A. Bell et al., 2011 Subspecific origin and haplotype diversity in the laboratory mouse. Nat Genet. 43: 648–655.

